# The Effect of Physical Activity Level on Age-Related Differences in Responses to Optic Flow Perturbation during Human Walking

**DOI:** 10.1101/2025.10.27.684797

**Authors:** Chunchun Wu, Tom J.W. Buurke, Rob den Otter, Claudine J.C. Lamoth, Menno P. Veldman

**Affiliations:** University of Groningen, University Medical Center Groningen, Department of Human Movement Sciences, Groningen, The Netherlands

**Keywords:** optic flow perturbations, walking, sensory reweighting, frequency-based analysis

## Abstract

**Background:** Human aging increases the reliance on vision for walking balance due to age-related declines in proprioceptive and vestibular function. Regular physical activity (PA) may reduce the reliance on visual input during walking. This study examined whether PA levels modulate age-related responses to perturbations of the optic flow that is crucial in the control of human locomotion.

**Methods:** Sixty active and inactive younger (YA and YI: 23.3±3.91 y) and older adults (OA and OI: 68.3±3.98 y; n=15 for each group) walked on a treadmill in front of a virtual hallway. The walking protocol consisted of 3-minute walking without, and 8-minute with mediolateral optic flow perturbation. Sacrum and heel marker positions and ground reaction forces were recorded. Power spectral density (PSD) of the mediolateral sacrum position and gait parameters were analyzed.

**Results:** The PSD increased more in OA compared to OI adults (p=0.041) while YA and YI adults did not differ. Mean (and variability of) step width and mediolateral margin of stability increased irrespective of age and PA (all p<0.001). During the 8-minute perturbation, OA adults demonstrated greater decreases in PSD than the OI adults (p=0.039). Additionally, the variability in the mediolateral margin of stability reduced more in YA and OA adults compared to YI and OI adults (p=0.048).

**Conclusion:** Higher PA levels in OA adults were associated with stronger immediate responses in body sway to optic flow perturbations compared to older inactive and younger adults. This may support the beneficial effects of physical activity on age-related visual dependency during gait.

## Introduction

Mobility is considered crucial for the quality of life and key in maintaining an independent lifestyle. Indeed, the EuroQol questionnaire [1] lists mobility as one of five determinants of quality of life. Mobility has been shown to depend on balance during walking [2], which is challenged with advancing age due to age-related declines in sensory systems, including the visual, proprioceptive, and vestibular systems [3]. For instance, reduced proprioceptive and vestibular function in older vs. younger adults is demonstrated by the compromised ability to detect changes in body position and movement during an upright stance [4] and a decreased vestibulo-ocular reflex [5]. These age-related declines consequently result in difficulties in balance control responses to changes in environmental demands, such as changes in surface conditions or walking speed [6]. Because visual function is relatively unaffected by age, older individuals rely on visual information for walking balance through sensory reweighting, a process defined as the shift to the use of reliable sources of sensory information when one or multiple sensory modalities are compromised [7,8].

Regular physical activity (PA) may reduce the dependency on visual information in older adults by protecting proprioceptive and vestibular functions [9,10,11]. For example, physical activity attenuates age-related reductions in proprioceptive function measured with a knee joint reproduction test [9] and active older adults showed superior vestibular function as compared to their sedentary counterparts, as indicated by 28-56% higher vestibular reflectivity [10]. Additionally, active older adults scored ∼30% higher than inactive older adults on the sensory organization test, which required them to maintain balance under demanding sensory conditions, suggestive of PA-related enhancement of sensory integration [11]. The PA-related effects on sensory function provide a rationale for the hypothesis that PA can positively affect balance. While an earlier study showed this for standing balance already [12], it is yet unclear whether these effects extend to walking balance and gait adaptability, which requires more complex sensory integration and dynamic regulation than postural control tasks [13].

The dependency on visual information for walking balance is often studied using optic flow perturbation. Initial responses to manipulations of the optic flow, i.e., making visual information unreliable, provide insights into the reliance on visual information for walking balance. Moreover, prolonged exposure to such perturbation provides insights into human’s ability to reweigh visual information when unreliable. Previous studies showed that perturbing the optic flow in the mediolateral direction during walking immediately (comparing baseline and early phases of perturbation) increased the step width (14%-20%) and mediolateral margins of stability (MoS, 6%), a measure of dynamic gait stability [14,15,16]. Interestingly, older adults show stronger immediate responses to mediolateral optic flow perturbations than young adults, as indicated by 4% and 5% higher step width and mediolateral MoS and a substantial 115-130% increase in step width variability [16,17,18]. Data showing that perturbation-induced body sway coincided with larger step widths suggest that changes in spatiotemporal gait parameters may be a direct consequence of affected trunk stability [15]. Moreover, follow-up studies indicated that the body sway was specific for the perturbation signal, as quantified by the power spectral density (PSD) of the mediolateral sacrum marker position. Larger perturbation-induced body sway in older individuals compared to young individuals (4-21 times larger; [19]) indeed suggest that older individuals rely more on visual information during optic flow perturbation than young adults. Following prolonged exposure to optic flow perturbation, adaption, defined as the change in walking parameters from early to late stages of the perturbation, was shown in both younger and older adults. Specifically, 7.3% larger step lengths and 17.4% narrower step widths during late versus early perturbation in older adults [17] were numerically similar in younger adults. Whether higher PA levels impact the ability to reweigh sensory information during walking under optic flow perturbations is yet unknown.

This study aimed to investigate whether the age-related response to immediate and prolonged exposure to optic flow perturbations is modulated by physical activity. Based on the age-related dependency on visual information, we hypothesized that (1) older adults show stronger immediate responses to optic flow perturbations than young adults and that PA attenuates these age-related responses. Similarly, we hypothesized that (2) older adults show reduced adaptability to optic flow perturbations compared to young adults and that PA would reduce these age-related declines in adaptability.

## Methods

### Participants

Healthy participants (N = 60) were divided into four groups: young active (YA, N=15), young inactive (YI, N=15), older active (OA, N=15), and older inactive (OI, N=15). Young and older adults were aged 18–30 and 60-75 years, respectively. The International Physical Activity Questionnaire (IPAQ) [20], which assesses the duration of leisure-time physical activity per week, was used to classify the participants as active or inactive. Participants were considered active and inactive when engaging in more or less than 300 minutes of total physical activity per week [21], respectively. Physical activity was calculated as the sum of the time spent on moderate-intensity activity and the time spent on vigorous-intensity activity multiplied by two. All participants were free of neurological, cognitive, or orthopedic impairments and had no previous experience with optic flow perturbation experiments. Written informed consent was provided before participating. The study protocol was approved by the Central Ethics Review Board of the University Medical Center Groningen (10543) and conducted according to the Declaration of Helsinki [22].

### Walking protocol

The protocol consisted of a continuous 14-minute walking session on an instrumented treadmill (M-Gait; Motek Medical B.V., Amsterdam, Netherlands). Walking conditions consisted of three phases: a 3-minute baseline phase, an 8-minute perturbation phase, and a 3-minute post-perturbation phase. The last 60 steps of the baseline phase were analyzed as baseline (BL). During the perturbation phase, the first 60 steps were considered early perturbation (EP) and the last 60 steps were considered late perturbation (LP). The post-perturbation phase helped participants acclimate before stepping off the treadmill but was not included in the analysis.(Figure 1A). During the experiment, participants walked through a virtual hallway displayed on a 180° semi-curved screen in front of the treadmill, which moved at a belt-speed-matched velocity in the anterior-posterior direction, programmed using D-Flow software (version 3.36.3, Motekforce Link, Amsterdam, Netherlands, Figure 1B). During the perturbation phase, mediolateral optic flow perturbations were superimposed on the baseline. Similar to previous studies, a perturbation signal P was generated [e.g., φ = 0, Equation 1, Figure 1A] and comprised the sum of three sinusoids.

**Figure 1.**
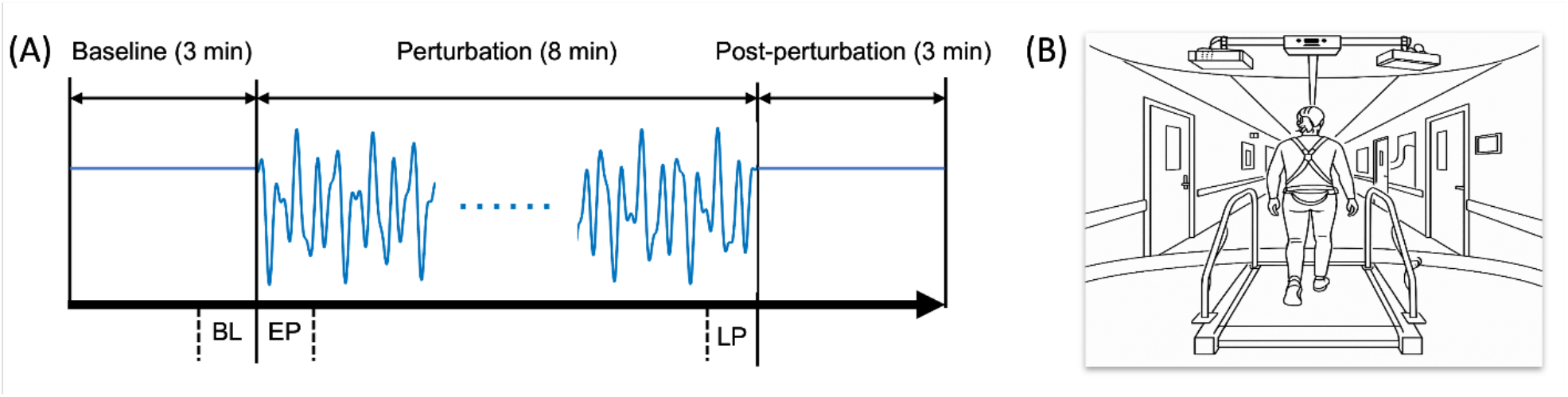
Walking protocol. (A) Walking protocol, with the duration of walking on the treadmill during each phase as well as the perturbation signal for the perturbation phase; (B) Participant walking on a treadmill in front of a virtual hallway. BL: the last 60 steps of baseline; EP: the first 60 steps of perturbation; LP: the last 60 steps of perturbation.

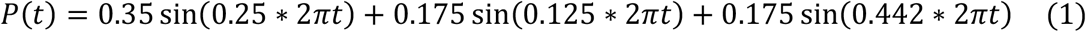

In our previous study examining the effect of gait speed on responses to optic flow perturbation, we evaluated three speeds (0.6, 1.2 and 1.8 m/s) [23], and showed that the effect of optic flow perturbations was most pronounced at the slowest speeds. Therefore, the current study focuses on 0.6 m/s for young and 0.5 m/s for older adults with a slight difference between young and older adults to mitigate age-related differences in fatigability. Walking speed was normalized to leg length by multiplying the gait speed by the square root of the leg length, quantified as the distance from the spina iliaca anterior superior to the medial malleolus of the right leg [24]. Before the walking protocol began, participants were given one minute to acclimate to the treadmill.

### Data acquisition

During the experiment, a 10-camera motion capture system (VICON MX, Vicon Motion Systems Ltd., Oxford, UK) was used to record the three-dimensional position data of the reflective markers placed on the sacrum, located midway between the posterior superior iliac spines and heels, located on the left and right calcaneus, at a sample frequency of 100 Hz. Two force plates embedded in the treadmill were used to collect ground reaction force data in six degrees of freedom, including 3D forces (Fx, Fy, Fz) and 3D moments (Mx, My, Mz), at a sample frequency of 1000 Hz. For safety purposes, participants were attached to a safety harness that did not provide body weight support.

### Data analysis

#### Pre-processing

Gaps in marker trajectories <25 frames (i.e., 250 ms) were interpolated with Woltring gap fill in Vicon Nexus (V2.12, Oxford Metrics Groups, Oxford, UK). Subsequently, a zero-phase low-pass Butterworth filter (2nd order, 15 Hz cutoff) was applied using custom software developed in MATLAB (vR2022B; The MathWorks Inc., Natick, MA, USA) to remove high-frequency noise while preserving the temporal characteristics of gait-related signals. The moments of heel strikes and toe-offs were determined by detecting the peaks and troughs in the absolute horizontal distance between the sacrum marker and heel markers, respectively [24].

The preprocessing of sacrum marker data for calculating the Power Spectral Density (PSD) was based on previous literature [19]. First, linear trends were removed from the raw signal using a detrending function. Next, a fourth-order, zero-lag low-pass Butterworth filter with a 12 Hz cut-off frequency was applied to attenuate high-frequency noise and ensure stable spectral estimation. Finally, the signal was normalized by dividing each value by the square root of the sum of the squared signal values [25].

#### Outcome parameters

To evaluate the degree to which the responses to the optic flow perturbations were related to the content of the perturbation signal, we quantified PSD of the sacrum marker position in the mediolateral direction at 0.250 Hz. This frequency was dominant in the perturbation signal (see Equation 1) and was therefore assumed to have the most impact on walking balance. To do so, pre-processed sacrum marker data were used to calculate the PSD using the ‘pwelch’ function implemented in MATLAB with a frequency resolution of 0.001 Hz. For, the BL, EP and LP phases the PSD was averaged around the dominant frequency of the perturbation signal with a 0.1 Hz tail.

The average step width and mediolateral MoS as well as their standard deviations, as an index of variability, were calculated over 60 steps for BL, EP and LP (see Figure 1 and walking protocol). Step width was calculated as the absolute mediolateral distance between the two heel markers at the time of consecutive heel strikes (i.e., one left and one right). The mediolateral MoS was computed based on the center of mass (CoM) position, which was estimated by integrating the ground reaction force-derived acceleration twice, with a high-pass filter (2nd-order Butterworth, 0.2 Hz) applied after the first integration to remove drift (Equation 2 [26]):

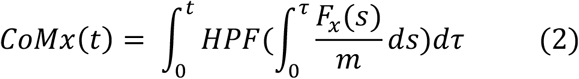

in which *F* is the mediolateral ground reaction force, and *m* is the weight of participants, HPF is High-Pass Filter. The extrapolated center of mass (XCoM) was derived from the CoM position and velocity by high-pass filtering (2nd-order Butterworth, 0.2 Hz) the CoM and low-pass filtering (2nd-order Butterworth, 0.2 Hz) the center of pressure (Equation 3 [26]):

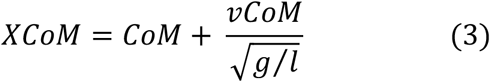

in which *vCoM* is mediolateral CoM velocity, *g* is gravitational acceleration and *l* is effective pendulum length defined as the distance between the spina iliaca anterior superior and the malleolus medialis. The mediolateral MoS was then determined as the minimum difference between the mean center of pressure (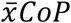) during a single stance and the XCoM (Equation 4 [27,28]):

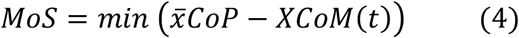

### Statistical analysis

IBM SPSS Statistics (version 28.0.1) was used to perform the statistical analyses. To test the hypotheses that immediate responses to optic flow perturbation and adaptation to prolonged exposure to optic flow perturbation were age- and PA-dependent, we performed two repeated measures analyses of variance (RM-ANOVAs) with two between-subject factors: PA (active vs. inactive) and Age (young vs. old) and one within-subject factor (Phase). To examine whether immediate responses differed between Age and PA, we performed a two (Phase: BL vs. EP) by two (Age) by two (PA) RM-ANOVA. To examine whether adaptability differed between Age and PA, we performed another two (Phase: EP vs. LP) by two (Age) by two (PA) RM-ANOVA. Greenhouse-Geisser correction was applied if this assumption was violated, and partial eta squared was used as a measure of the effect size. A significance level of p < 0.05 was used for all statistical tests.

## Results

Descriptive statistics of the participants’ characteristics, anthropometric data, and physical activity levels are presented in Table 1.

**Table 1.** Basic information, anthropometric data and physical activity levels of participants.

|  |  |  |  |  |  |  |
| --- | --- | --- | --- | --- | --- | --- |
|  | Young active | Young inactive | Older active | Older inactive | F | p |
| Gender (F/M) | 8/7 | 8/7 | 8/7 | 7/8 | F (3,56) = 0.062 | 0.979 |
| Age (years) | 22.73 ± 3.81 | 23.93 ± 4.04 | 68.73 ± 4.08 | 67.87 ± 3.98 | F (3,56) = 638.97 | < 0.001 |
| Height (cm) | 179.98 ± 10.34 | 175.66 ± 11.32 | 173.25 ± 7.63 | 174.95 ± 7.72 | F (3,56) = 1.39 | 0.254 |
| Weight (kg) | 76.53 ± 13.09 | 71.65 ± 17.79 | 74.97 ± 16.93 | 73.12 ± 10.58 | F (3,56) = 0.307 | 0.820 |
| Leg (cm) | 92.97 ± 6.84 | 90.07 ± 6.77 | 87.49 ± 11.18 | 90.13 ± 4.66 | F (3,56) = 1.25 | 0.299 |
| Physical activity (min/week) | 751.33 ± 311.40 | 82.33 ± 87.52 | 681.33 ± 351.71 | 78.53 ± 92.71 | F (3,56) = 34.34 | < 0.001 |

### Immediate responses to optic flow perturbation in all parameters

As illustrated in representative participants in Figure 2, PSD of the sacrum marker increased by 1017.4% across all groups from baseline to early perturbation (p < 0.001, Table S1). On a group level, Figure 3 shows a significant Age by PA by Phase interaction indicating that PSD increased more in OA adults than in OI adults (p = 0.041, Figure 3A). This finding was further supported by the Age by PA interaction in PSD (p = 0.026, Table S1) revealing that OA adults showed higher PSD than OI adults, while YA and YI adults did not differ.

**Figure 2.**
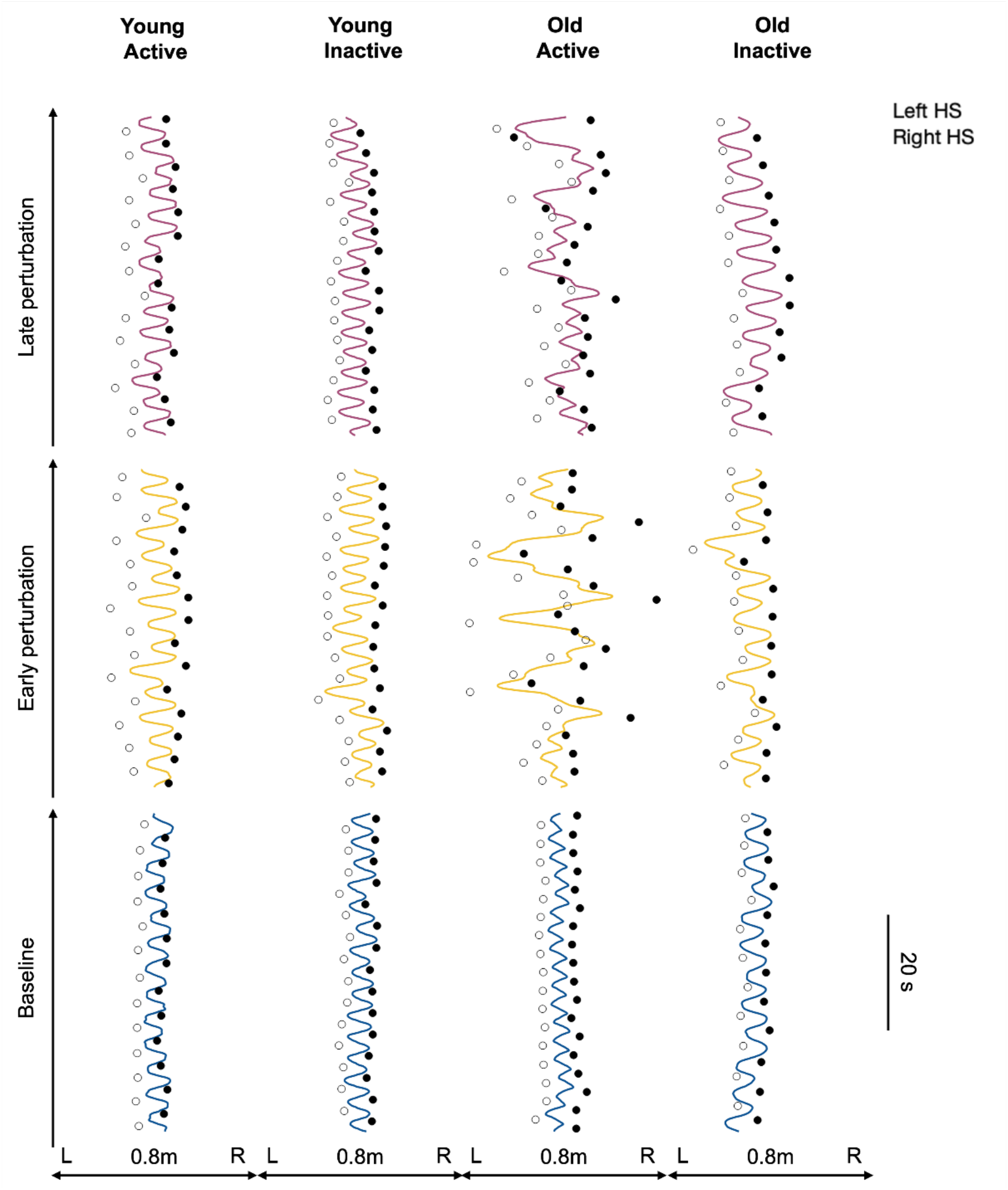
Mediolateral sacrum position trajectories and foot placements across groups and phases. The figure illustrates the mediolateral sacrum trajectories and left (open circles) and right (filled circles) foot placements during baseline (blue), early perturbation (yellow) and late perturbation (purple) in a representative participant from each group. HS: heel strike.

**Figure 3.**
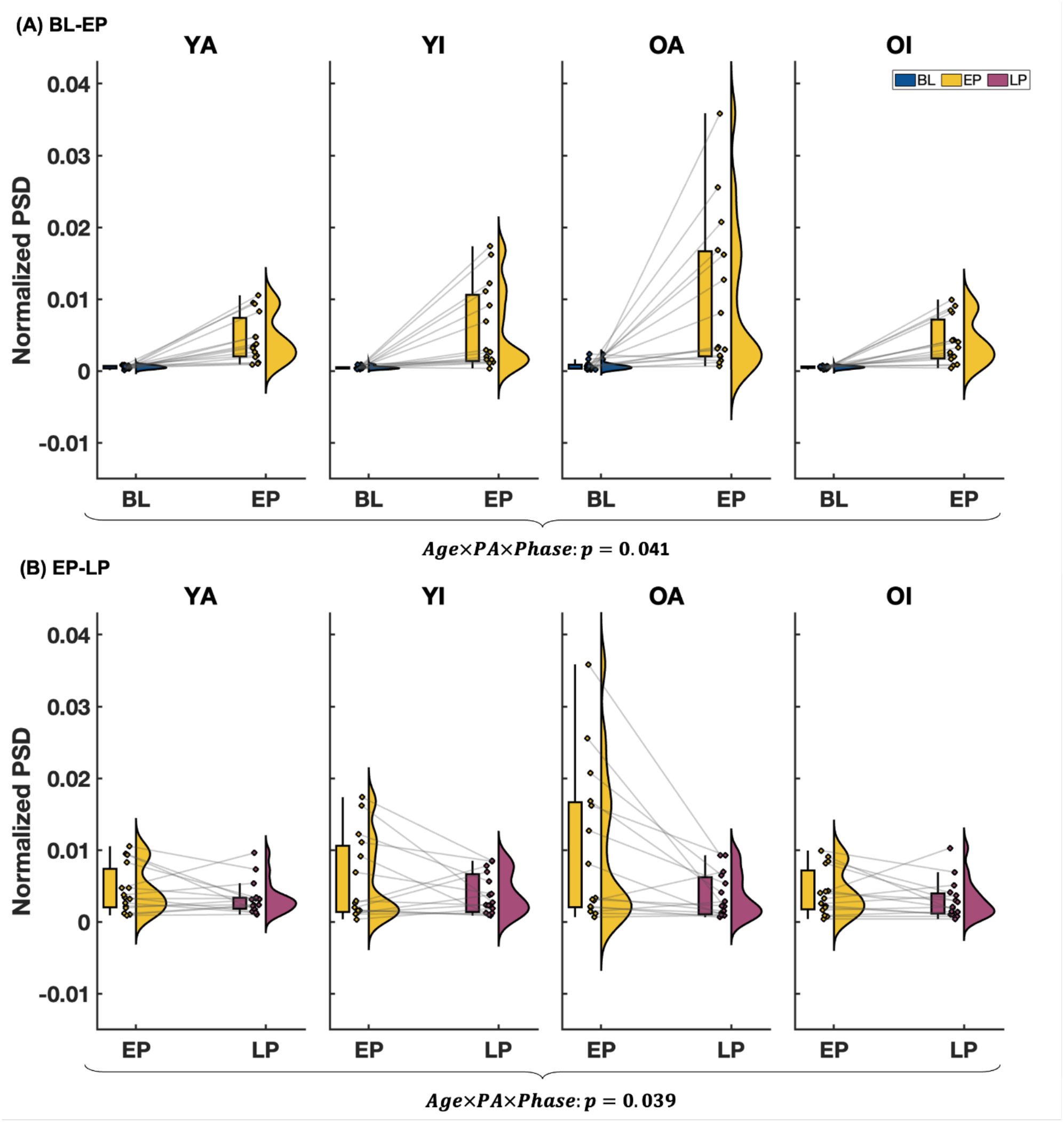
Response of normalized power spectral density to optic flow perturbations over phases. (A) Comparison between baseline and early perturbation; (B) Comparison between early perturbation and late perturbation; PSD: power spectral density; OA: older active adults; OI: older inactive adults; YA: young active adults; YI: young inactive adults; BL: baseline; EP: early perturbation; LP: late perturbation. Bars indicate group means ± standard deviation, the dots and lines indicate individual data and within-subjects changes, the semi-violin plots indicate the overall data distribution.

As shown in Figure 4, optic flow perturbation increased step width (38.6%), mediolateral MoS (54.9%) and their variabilities (172.1% and 257.7%, respectively) independent of Age and PA (all p < 0.001, Table S1). In addition, no significant interactions between Phase and Age or PA for step width, mediolateral MoS, and their variabilities were observed.

**Figure 4.**
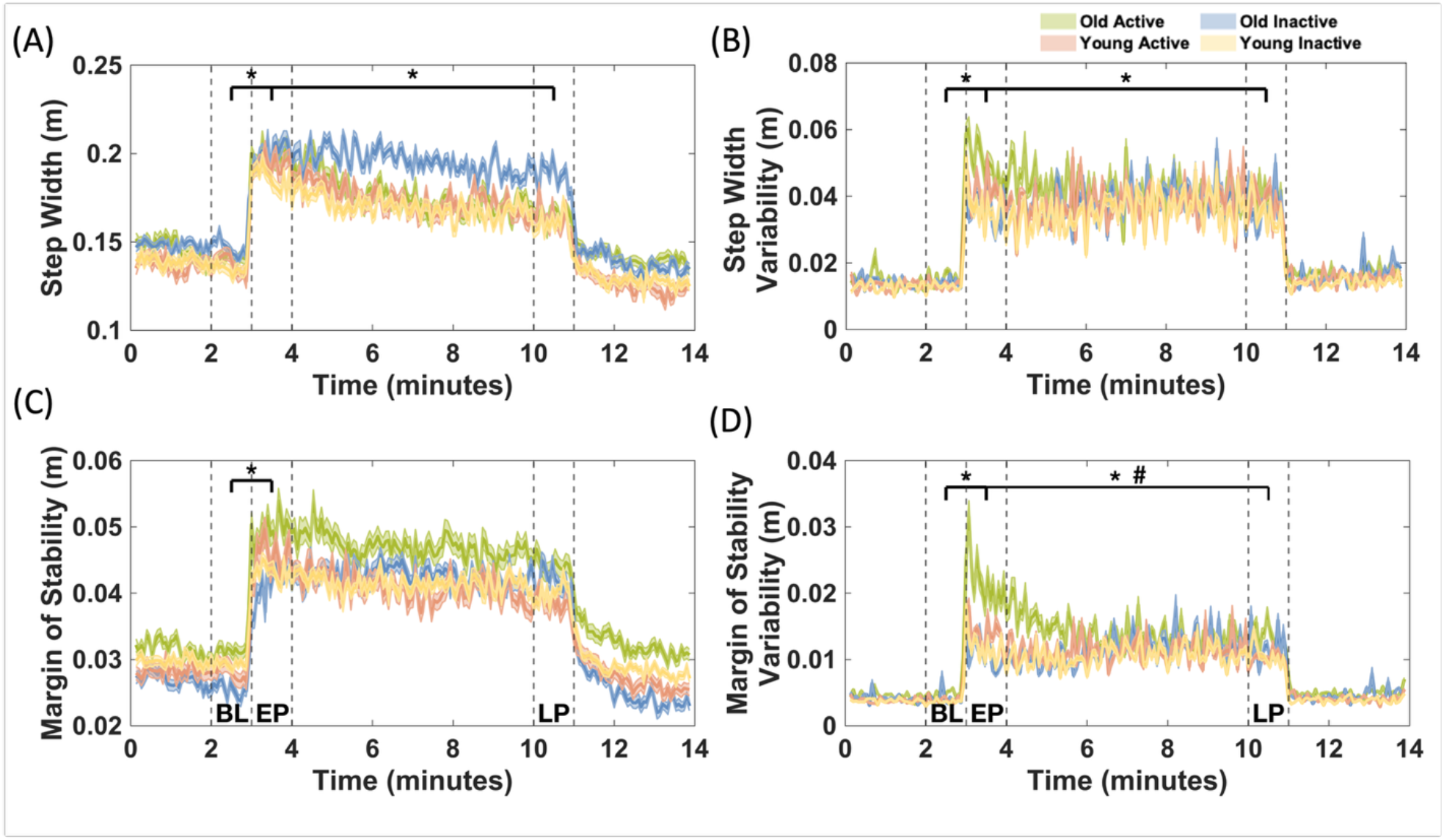
Changes in kinematic gait parameters in four groups during exposure to optic flow perturbations. The figure shows the mean ± standard error of (A) Step Width, (B) Step Width Variability, (C) Mediolateral Margin of Stability, and (D) Mediolateral Margin of Stability Variability. The parameters are averaged across consecutive 4-second bins to improve visibility. BL: baseline; EP: early perturbation; LP: late perturbation. ^*^: main effects of Phase (all p < 0.01); #: Physical Activity by Phase interaction (p = 0.048).

### Adaptation to the optic flow perturbation

Overall, the participants changed their responses following prolonged exposure to the perturbation. PSD decreased from early to late perturbation (p < 0.005, Table S2, Figure 2). Figure 3B illustrates that the decrease in PSD was greater in OA than in OI, YA and YI from EP to LP, as reflected by an Age by PA by Phase interaction (p = 0.039, Table S2, Figure 3B). This effect drove the Age by PA interaction (p = 0.034, Table S2) indicating OA showed larger PSD than OI adults, and YA adults showed smaller PSD than YI adults.

Figure 4 shows the changes in step width, mediolateral margin of stability and their variabilities over the course of the walking protocol. From EP to LP, the mean and variability of step width significantly decreased with 11.8% and 10.9% respectively, as well as the variability in mediolateral MoS (29.0%, all p<0.005, Table S2). Moreover, a significant PA by Phase interaction indicated that active individuals showed a greater reduction in the variability of mediolateral MoS compared to inactive adults (p = 0.048, Table S2, Figure 4).

## Discussion

This study examined whether the age-related responses to immediate and prolonged exposure to optic flow perturbations were modulated by physical activity levels. Results indicated that in contrast with our hypothesis the immediate effects of optic flow perturbation on mean and variability of step width and mediolateral MoS were independent of age and PA. In addition, the increases in PSD from BL to EP were larger in older active compared to older inactive adults and young active and inactive adults. With prolonged perturbations, older active adults showed a larger reduction in PSD and mediolateral MoS variability than inactive older adults. Together, the present results may support the beneficial effects of physical activity on age-related responses to visual perturbations during gait.

Because the changes in PSD from BL to EP were Age- and PA-dependent while the increases in (the variability of) step width and mediolateral MoS were not, changes in body sway appear to be more sensitive to optic flow perturbations, and therefore might represent a more direct consequence of visual perturbations. The larger increase in body sway in active older adults could indicate a more optimal use of degrees of freedom compared to inactive older adults. Specifically, a previous study has shown a relationship between available degrees of freedom and flexibility in balance recovery strategies (e.g., [29]). The results of the present study may indicate that in older adults, who are arguably more susceptible to visual perturbations compared to younger adults due to increased reliance on visual information for walking balance [19], a physically active lifestyle allowed to exploit more degrees of freedom for a more complex response to maintain balance compared to inactive older adults. Although the finding that inactive older adults did not increase their body sway in response to optic flow perturbation was unexpected, it is possible that they rather reduced their degrees of freedom in response to perturbation, thereby limiting the complexity of control [30]. Although speculative, this interpretation agrees with studies showing reduced movement complexity in conditions with impaired neuromuscular function [31] and reduced ability to adapt to perturbations in conditions that affect proprioception [32]. However, whether or not this speculation extends to degrees of freedom related to body sway needs to be explored in future studies. Moreover, higher PA levels in older adults have been associated with preserved neuromuscular control and muscle mechanical properties [33] and enhanced sensorimotor integration, enabling efficient detection and adaptation to dynamic environmental changes [34]. If true, this may not only explain the flexible and complex responses in the older active group on average, but also the variability in the change in body sway in response to the onset of optic flow perturbation within this group. That is, the IPAQ questionnaire does not distinguish between different types of PA, whereas it is possible that the type of PA influences neuromuscular control and therefore the response to optic flow perturbations.

At the end of the perturbation phase (LP), the magnitude of body sway and (the variability in) step width and mediolateral MoS margin of stability was similar between groups. While this indicates overall adaptation towards baseline walking behavior, differences between groups in initial responses to the optic flow perturbation, i.e., from BL to EP, complicate the comparison of the adaptation phase between groups and therefore the interpretation of the underlying mechanisms. Adaptation to optic flow perturbation may have resulted from sensory reweighting, the gradual shift to reliance on different sensory modalities when a given modality becomes unreliable [7,8]. Through this process, the reliance on visual information decreases and as a result, the magnitude of body sway decreased. Since a previous study showed a relationship between reductions in body sway and decreases in hip abduction angles following sudden perturbations that challenge trunk stability [35], it would be interesting exploit 3D kinematic analyses to evaluation changes in joint angles in response to optic flow perturbations. Furthermore, the perturbation magnitude could be personalized in future research to match initial responses between groups, enabling a valid comparison of the adaptation process, similar to a previous approach normalizing stability in different walking speeds [36].

A limitation of the present study was the fact that eye-movements were not measured, which could have provided insights into alternative visual strategies independent of the body movements that this study focused on. That is, participants may have used specific fixations or saccades to maintain balance. For example, earlier research showed that humans’ gaze behavior aligns with changes in visual information [37]. Incorporating eye-tracking may be an interesting follow-up in the context of age- and PA-related responses to optic flow perturbations to quantify fixations and saccades during the perturbation phase to obtain insights into the use of visual information while walking under optic flow perturbations.

## Conclusion

This study demonstrates that PA levels influence responses to optic flow perturbation differently in younger and older adults. Altogether, the present results may support the beneficial effects of physical activity on age-related responses to gait perturbations.

## CRediT authorship contribution statement

Chunchun Wu: Writing – review & editing, Writing – original draft, Investigation, Visualization, Methodology, Conceptualization, Data analysis. Tom J.W. Buurke: Writing – review & editing, Visualization, Methodology, Conceptualization, Data analysis. Rob den Otter: Writing – review & editing, Methodology, Conceptualization. Claudine J.C. Lamoth: Writing – review & editing, Methodology, Conceptualization. Menno P. Veldman: Writing – review & editing, Writing – original draft, Visualization, Methodology, Conceptualization, Data analysis.

## Funding and acknowledgement

This work was supported by China Scholarship Council – University of Groningen Scholarship (C. Wu: 202206320033).

## Data availability

Due to data protection, the datasets collected and analyzed during this study are not openly available. However, reasonable requests for data supporting the findings of this study are available from the corresponding author upon reasonable request.

