## Supplementary table for "The Effect of Physical Activity Level on Age-Related Differences in Responses to Optic Flow Perturbation during Human Walking"

Table S1: Effects of age, and physical activity on immediate adjustment to optic flow perturbation

| | Young active | | Young inactive | | Old active | | Old inactive | | Statistics | | | |
|---|---|---|---|---|---|---|---|---|---|---|---|---|
| | BL | EP | BL | EP | BL | EP | BL | EP | Effect | F | P | Partial $\eta^2$ |
| SW | 0.137±0.050 | 0.192±0.061 | 0.134±0.044 | 0.187±0.054 | 0.153±0.043 | 0.196±0.060 | 0.141±0.043 | 0.198±0.059 | Age | F(1,56)=0.002 | 0.535 | 0.007 |
| | | | | | | | | | PA | F(1,56)=0.001 | 0.981 | 0.000 |
| | | | | | | | | | Phase | F(1,56)=134.555 | <0.001 | 0.706 |
| | | | | | | | | | Age*PA | F(1,56)=0.090 | 0.766 | 0.002 |
| | | | | | | | | | Age*phase | F(1,56)=0.009 | 0.927 | 0.000 |
| | | | | | | | | | PA*phase | F(1, 56)=0.104 | 0.748 | 0.002 |
| | | | | | | | | | Age*PA*phase | F(1,56)=0.018 | 0.893 | 0.000 |
| MoS | 0.028±0.015 | 0.045±0.024 | 0.029±0.008 | 0.043±0.011 | 0.031±0.013 | 0.049±0.022 | 0.025±0.011 | 0.040±0.018 | Age | F(1,56)=0.000 | 0.998 | 0.000 |
| | | | | | | | | | PA | F(1,56)=0.847 | 0.361 | 0.015 |
| | | | | | | | | | Phase | F(1,56)=111.364 | <0.001 | 0.665 |
| | | | | | | | | | Age*PA | F(1,56)=0.763 | 0.386 | 0.013 |
| | | | | | | | | | Age*phase | F(1,56)=0.036 | 0.849 | 0.001 |
| | | | | | | | | | PA*phase | F(1,56)=0.804 | 0.374 | 0.014 |
| | | | | | | | | | Age*PA*phase | F(1,56)=0.001 | 0.970 | 0.000 |
| SWV | 0.017±0.004 | 0.046±0.018 | 0.016±0.003 | 0.041±0.024 | 0.018±0.009 | 0.056±0.035 | 0.018±0.004 | 0.043±0.021 | Age | F(1,56)=1.455 | 0.233 | 0.025 |
| | | | | | | | | | PA | F(1,56)=1.819 | 0.183 | 0.031 |
| | | | | | | | | | Phase | F(1,56)=83.253 | <0.001 | 0.598 |
| | | | | | | | | | Age*PA | F(1,56)=0.239 | 0.627 | 0.004 |
| | | | | | | | | | Age*phase | F(1,56)=0.475 | 0.493 | 0.008 |
| | | | | | | | | | PA*phase | F(1,56)=1.539 | 0.220 | 0.027 |
| | | | | | | | | | Age*PA*phase | F(1,56)=0.392 | 0.534 | 0.007 |
| MoSV | 0.004±0.001 | 0.016±0.008 | 0.004±0.001 | 0.013±0.009 | 0.005±0.002 | 0.025±0.023 | 0.005±0.003 | 0.015±0.010 | Age | F(1,56)=2.948 | 0.091 | 0.050 |
| | | | | | | | | | PA | F(1,56)=3.255 | 0.077 | 0.055 |
| | | | | | | | | | Phase | F(1,56)=50.339 | <0.001 | 0.473 |
| | | | | | | | | | Age*PA | F(1,56)=1.044 | 0.311 | 0.018 |
| | | | | | | | | | Age*phase | F(1,56)=1.431 | 0.237 | 0.025 |
| | | | | | | | | | PA*phase | F(1,56)=3.149 | 0.081 | 0.053 |
| | | | | | | | | | Age*PA*phase | F(1,56)=1.298 | 0.260 | 0.023 |
| PSD | 0.531±0.215 | 4.501±3.316 | 0.467±0.205 | 5.851±5.885 | 0.716±0.593 | 10.191±10.775 | 0.495±0.162 | 4.134±3.230 | Age | F(1,56)= 1.597 | 0.212 | 0.028 |
| | | | | | | | | | PA | F(1,56)= 2.272 | 0.137 | 0.039 |
| | | | | | | | | | Phase | F(1,56)= 41.911 | <0.001 | 0.428 |
| | | | | | | | | | Age*PA | F(1,56)= 5.216 | 0.026 | 0.085 |
| | | | | | | | | | Age*phase | F(1,56)= 1.174 | 0.283 | 0.021 |
| | | | | | | | | | PA*phase | F(1,56)= 1.623 | 0.208 | 0.028 |
| | | | | | | | | | Age*PA*phase | F(1,56)= 4.362 | 0.041 | 0.072 |

Table S2: Effects of age, and physical activity on prolonged adjustment to optic flow perturbation

| | Young active | | Young inactive | | Old active | | Old inactive | | Statistics | | | |
|---|---|---|---|---|---|---|---|---|---|---|---|---|
| | EP | LP | EP | LP | EP | LP | EP | P | Effect | F | P | Partial η2 |
| SW | 0.192±0.061 | 0.163±0.053 | 0.187±0.054 | 0.163±0.042 | 0.196±0.060 | 0.166±0.045 | 0.198±0.059 | 0.189±0.048 | Age | F(1,56)=0.755 | 0.389 | 0.013 |
| | | | | | | | | | PA | F(1,56)=0.130 | 0.719 | 0.002 |
| | | | | | | | | | Phase | F(1,56)=22.789 | <0.001 | 0.289 |
| | | | | | | | | | Age*PA | F(1,56)=0.326 | 0.570 | 0.006 |
| | | | | | | | | | Age*phase | F(1,56)=0.607 | 0.439 | 0.011 |
| | | | | | | | | | PA*phase | F(1,56)=1.750 | 0.191 | 0.030 |
| | | | | | | | | | Age*PA*phase | F(1,56)=0.752 | 0.369 | 0.013 |
| MoS | 0.045±0.024 | 0.039±0.018 | 0.043±0.011 | 0.040±0.010 | 0.049±0.022 | 0.044±0.019 | 0.040±0.018 | 0.042±0.014 | Age | F(1,56)=0.183 | 0.670 | 0.003 |
| | | | | | | | | | PA | F(1,56)=0.345 | 0.559 | 0.006 |
| | | | | | | | | | Phase | F(1,56)=3.598 | 0.063 | 0.060 |
| | | | | | | | | | Age*PA | F(1,56)=0.388 | 0.536 | 0.007 |

| | | | | | | | | | | | |
|---|---|---|---|---|---|---|---|---|---|---|---|
| | | | | | | | | | Age*phase | F(1,56)=1.020 | 0.317 | 0.018 |
| | | | | | | | | | PA*phase | F(1,56)=2.508 | 0.119 | 0.043 |
| | | | | | | | | | Age*PA*phase | F(1,56)=0.213 | 0.647 | 0.004 |
| | | | | | | | | | Age | F(1,56)=0.481 | 0.491 | 0.009 |
| | | | | | | | | | PA | F(1,56)=1.093 | 0.300 | 0.019 |
| | | | | | | | | | Phase | F(1,56)=11.024 | 0.002 | 0.164 |
| SWV | 0.046±0.018 | 0.041±0.024 | 0.041±0.024 | 0.036±0.018 | 0.056±0.035 | 0.041±0.024 | 0.043±0.021 | 0.039±0.021 | Age*PA | F(1,56)=0.037 | 0.847 | 0.001 |
| | | | | | | | | | Age*phase | F(1,56)=1.375 | 0.246 | 0.024 |
| | | | | | | | | | PA*phase | F(1,56)=1.664 | 0.202 | 0.029 |
| | | | | | | | | | Age*PA*phase | F(1,56)=1.577 | 0.214 | 0.027 |
| | | | | | | | | | Age | F(1,56)=2.403 | 0.127 | 0.041 |
| | | | | | | | | | PA | F(1,56)=2.149 | 0.148 | 0.037 |
| | | | | | | | | | Phase | F(1,56)=14.165 | <0.001 | 0.202 |
| MoSV | 0.016±0.008 | 0.011±0.007 | 0.013±0.009 | 0.010±0.004 | 0.025±0.023 | 0.014±0.010 | 0.015±0.010 | 0.013±0.008 | Age*PA | F(1,56)=0.696 | 0.408 | 0.012 |
| | | | | | | | | | Age*phase | F(1,56)=0.967 | 0.330 | 0.017 |
| | | | | | | | | | PA*phase | F(1,56)=3.149 | 0.048 | 0.053 |
| | | | | | | | | | Age*PA*phase | F(1,56)=1.298 | 0.195 | 0.023 |
| | | | | | | | | | Age | F(1,56)=0.635 | 0.429 | 0.011 |
| | | | | | | | | | PA | F(1,56)= 1.260 | 0.266 | 0.022 |
| | | | | | | | | | Phase | F(1,56)= 12.842 | <0.001 | 0.187 |
| PSD | 4.501±3.316 | 3.393±2.362 | 5.851±5.885 | 3.915±2.828 | 10.191±10.775 | 3.667±3.070 | 4.134±3.230 | 3.067±2.685 | Age*PA | F(1,56)= 4.002 | 0.050 | 0.067 |
| | | | | | | | | | Age*phase | F(1,56)= 2.348 | 0.131 | 0.040 |
| | | | | | | | | | PA*phase | F(1,56)= 2.433 | 0.124 | 0.042 |
| | | | | | | | | | Age*PA*phase | F(1,56)= 4.484 | 0.039 | 0.074 |
